# Epigenetic loss of heterozygosity of *Apc* and an inflammation-associated mutational signature detected in *Lrig1*^+/-^-driven murine colonic adenomas

**DOI:** 10.1101/650879

**Authors:** Jessica Preston

## Abstract

Murine colonic adenomas induced by the loss of a single copy of the tumor suppressor gene *Apc* in *Lrig1*^*+/-*^ expressing progenitor cells grow rapidly, with high penetrance and tumor multiplicity. This study investigates the prevalence of intertumoral genetic heterogeneity and phenotypic variation across tumors, and attempts to identify the genomic cause of the unusual phenotype. Adult *Lrig1-CreERT2/+; Apc-flox/+* mice were intraperitoneal injected with 2mg tamoxifen for 3 consecutive days which induced highly penetrant, dysplastic adenomas in the distal colon ∼100 days later. Whole tumors (n=14) from 8 mice were excised and tumor exome DNA and mRNA were sequenced. Somatic mutations present in the tumor exome DNA were compared with adjacent normal colon (n=9 tumors from 3 mice). Putative somatic mutations were called after stringent filtering using SeuratSomatic, a Genome Analysis Toolkit software module. RNA-Seq was performed on tumor mRNA (n=5 tumors from 5 mice) compared to wildtype colon (n=3). Differential gene expression was profiled using the R package DESeq2. Copy number variations and splicing defects were assessed using custom tracks on the UCSC genome browser.

Adenomas resulting from the loss of *Apc* in *Lrig1*^*+/-*^*-* expressing colonic progenitor cells are genetically heterogeneous and hypermutated, containing ∼25-30 high-quality somatic mutations per megabase. A loss of heterozygosity of *Apc* was not observed in the tumor genomes, however evidence of an epigenetic loss of heterozygosity was readily apparent in the tumor transcriptome. The tumors display a strong bias toward G: C > A: T point mutations, which are a signature of guanine adducts, associated with tobacco tar and *H. pylori* infections. Putative tumor-driving mutations were detected and thousands of differentially expressed genes were identified including several UDP glucuronosyltransferases. Abnormal splicing patterns characterized by a loss of intron retention were detected in several RNA-binding genes throughout the tumor transcriptome. The widespread defects in gene expression, genomic stability, and splicing patterns implies that an early epigenetic loss of *Apc* in *Lrig1*^*+/-*^-expressing progenitor cells causes a rapid formation of guanine adducts and a corresponding accumulation of mutation C>A point mutations. This study demonstrates that randomly-appearing oncogenic mutations can become fixed into a latent genomic reservoir prior to advanced disease.

## Introduction

Human colorectal cancer (CRC) is the second leading cause of cancer death in the US, and ∼25% of CRC patients are incurable at the time of diagnosis ^1,2^. CRC rates are sharply increasing in younger patients, who tend to be diagnosed at an advanced stage when there are few effective targeted therapies available to treat the disease. Genetic heterogeneity is inherent to the disease, providing tumor cells with the ability to rapidly adapt and resist treatments ^3,4^. Currently CRC is clinically segregated into four broad subcategories based on the expression of various biomarker molecules called consensus molecular subtypes, however this classification system greatly oversimplifies the diversity and interrelatedness of cancer cell species ^5^. Recent breakthroughs in ‘tumor-omics’ research have revealed countless genes and growth mechanisms involved in regulating tumor pathology, but lists of biomarkers and knowledge about signaling pathways have not yet translated into substantial disease cures. The specific steps required to counteract the critical aspects of CRC progression remain very poorly understood despite decades of research. The difficulty in treating CRC is due to the inevitable unpredictability of natural selection, which enables cancer cells to behave as moving targets to evade treatment through constant evolution. Tumors often acquire resistance to cancer therapies by accumulating slight alterations in immune signaling, metabolic status, and ectopic gene expression via the accumulation of *de novo* somatic mutations, and the existence of several subtypes of CRC further complicates treatment since each subtype presents its own unique set of challenges.

The cell-of-origin of CRC derives from a population of rapidly-dividing colonic stem cells located at the base of the intestinal epithelial crypts, which are identifiable based on the expression of leucine-rich repeat containing G protein-coupled receptor 5 (*Lgr5*). *Lgr5* is the downstream target of *R-spondin* in the canonical *Wnt/ β-catenin* pathway. Mutations in the tumor suppressor gene Adenomatous Polyposis Coli (*Apc*) and other members of the canonical *Wnt* pathway are the hallmark of CRC ^6-9^; loss of heterozygosity (LOH) of *Apc* tends to occur during the early stages of CRC tumorigenesis and precipitates a cascade of deleterious effects which disrupt a cell’s ability to regulate the cell cycle.

Human tumors often exhibit distinct patterns of somatic mutations based on the specific category of mutation and the background genomic sequence context of the damage ^10-12^. These somatic signatures can vary greatly depending on the anatomical location and carcinogenic agents involved in a given tumor and can sometimes reveal sources of mutation. For example, tumors caused by tobacco tar contain an abundance of C > A transition point mutations due to the formation of guanine adducts. Our current understanding of the variation of somatic genomic landscapes across CRC cells and tumors within is a host is incomplete due to the highly heterogeneous nature of CRC ^5^. Functional DNA mismatch repair enzymes are required for maintaining genomic integrity, but the specific signaling mechanisms involved in regulating these processes remain unresolved ^13-16^.

Leucine-rich repeats and immunoglobulin-like domains 1 (*Lrig1)* is a transmembrane feedback regulator of growth factor receptor tyrosine kinases that is expressed in the *Lgr5*^+^ stem cell population present at the base of the colonic crypts ^17-20^. *Lrig1* acts as a tumor suppressor gene in several contexts ^21-26^. The *Lrig1-CreERT2/+; Apc-flox/+* inducible mouse model of colonic adenoma is based on the conditional Cre-recombinase-driven loss of a single copy of *Apc* under the control of the (*Lrig1*) promoter, referred to as the *Lrig1-(Apc-flox)* model hereafter. The colonic stem cells of the *Lrig1-(Apc-flox)* mice express one single wildtype copy each of the *Lrig1* and *Apc* genes after the engineered recombination in *Lrig1*^*+/-*^-expressing *Lgr5*^*+*^ stem cells. Within 100 days of the loss of one copy of *Apc*, rapidly growing adenomas with extremely high tumor penetrance and multiplicity appear in the distal colon ^27^.

Mouse models of colorectal cancer vary greatly in terms of tumor aggressiveness and resemblance to human tumors. Intriguingly, two independent mouse models of CRC tumorigenesis display uncanny similarity to each other in terms of tumor onset, penetrance, multiplicity, anatomical location, and mortality: *Lrig1-CreERT2/+; Apc-flox/+* model and the *Glutathione transferase(Gst)-null; Apc*^*Min*^ model. Glutathione transferases (*Gsts*) typically catalyze the conjugation of carcinogens to glutathione ^28^. Results from a study from 2009 which crossed a *Gst*-null allele into the standard *Apc*^Min^ murine mouse model of CRC and reported much earlier tumor onset, decreased survival, a 6-fold increase in distal colorectal adenoma incidence and a 50-fold increase in adenoma multiplicity compared to Apc^Min^ mice ^29^. The authors noted that the colons of these mice expressed higher levels of the inflammatory genes *IL-6, IL-4*, and nitric oxide synthase and concluded that *Gsts* regulate the inflammatory response of the colon and their loss rapidly accelerates the course of tumor progression. The remarkable phenotypic similarities between the *Lrig1-(Apc-flox)* and *Gst-null; Apc*^*Min*^ mouse models imply that a common mechanism of tumor formation is responsible in both cases.

This work seeks to understand the genomic changes occurring during the early stages of tumorigenesis in rapidly-growing *Lrig1-(Apc-flox)* murine colonic adenomas. Exomic DNA and mRNA captured from the *Lrig1-(Apc-flox)* adenomas was sequenced in order to detect the presence of transcriptomic and genomic alterations. Specifically, the genetic heterogeneity and somatic signatures of tumors from a single mouse were assessed using exome DNA profiling and the prevalence of differential gene expression and splicing defects was assessed across tumors using mRNA-Seq.

## Results

Adenomas resulting from the inducible loss of *Apc* in *Lrig1*^+/-^-expressing colonic epithelial cells are hypermutated and genetically heterogeneous. Tumor exome DNA contained 930-1300 high-quality somatic mutations per tumor (∼25-30 mutations per megabase) distributed uniformly throughout the genomie, with virtually identical frequencies of nucleotide motifs observed across all nine tumors sequenced (**Figure 1**). There were no large copy number variations or rearrangements detected. The adenomas exhibited an overwhelming abundance of an C to A transversion point mutations with remarkably similar frequencies. The somatic mutations displayed nearly identical patterns of background genomic sequence context across the nine tumors (**Figure 2**). This type of mutational signature is often seen in smoking-induced lung cancers and stomach cancers induced by *H. pylori* infection, which are associated with an increase in the formation of guanine adducts. The overwhelming abundance of C to A transversions observed in the tumor exomes implies that the loss of one copy of *Apc* in *Lrig1*^+/-^ colonic stem cells caused an increased mutation rate due to the inability of cells to repair DNA adducts caused by the oxidation of guanine.

**Figure 1.**
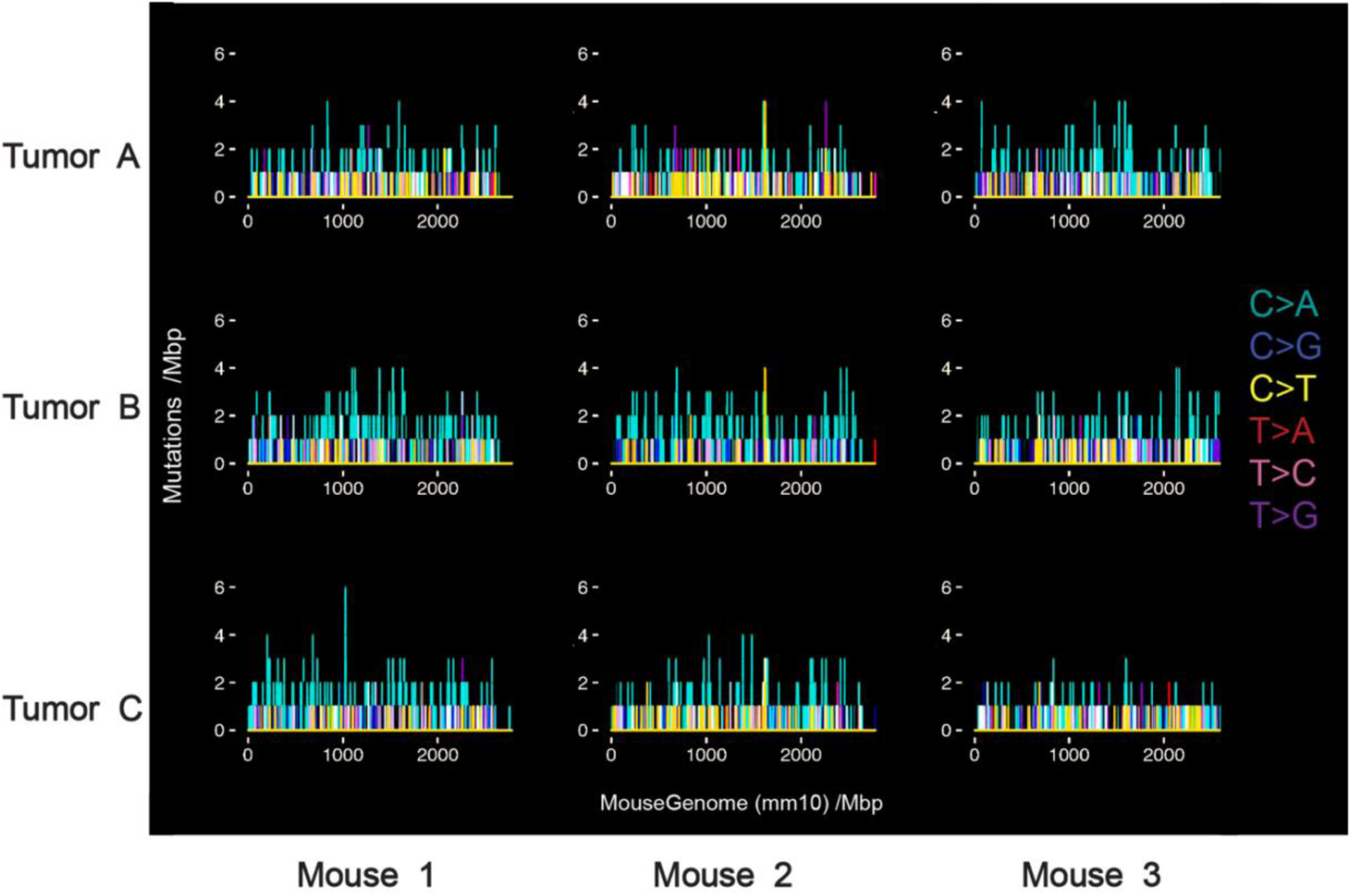
Colonic adenomas resulting from inducible loss of *APC* in Lrig1+/- stem cells are hypermutated and genetically heterogeneous. Exome sequencing of adenomas from *Lrig1-Cre/+;Apcfl/+* mice detected the presence of ∼25-35 high-quality somatic mutations per megabase, with an abundance of C:G>A:T transversions point mutations. The number and type of somatic mutations from each adenoma exome is plotted. There are 9 adenomas from 3 mice; each column plots three tumors from a single mouse. Similar mutational frequencies and motif distributions were observed across the nine tumor exomes sequenced.

**Figure 2.**
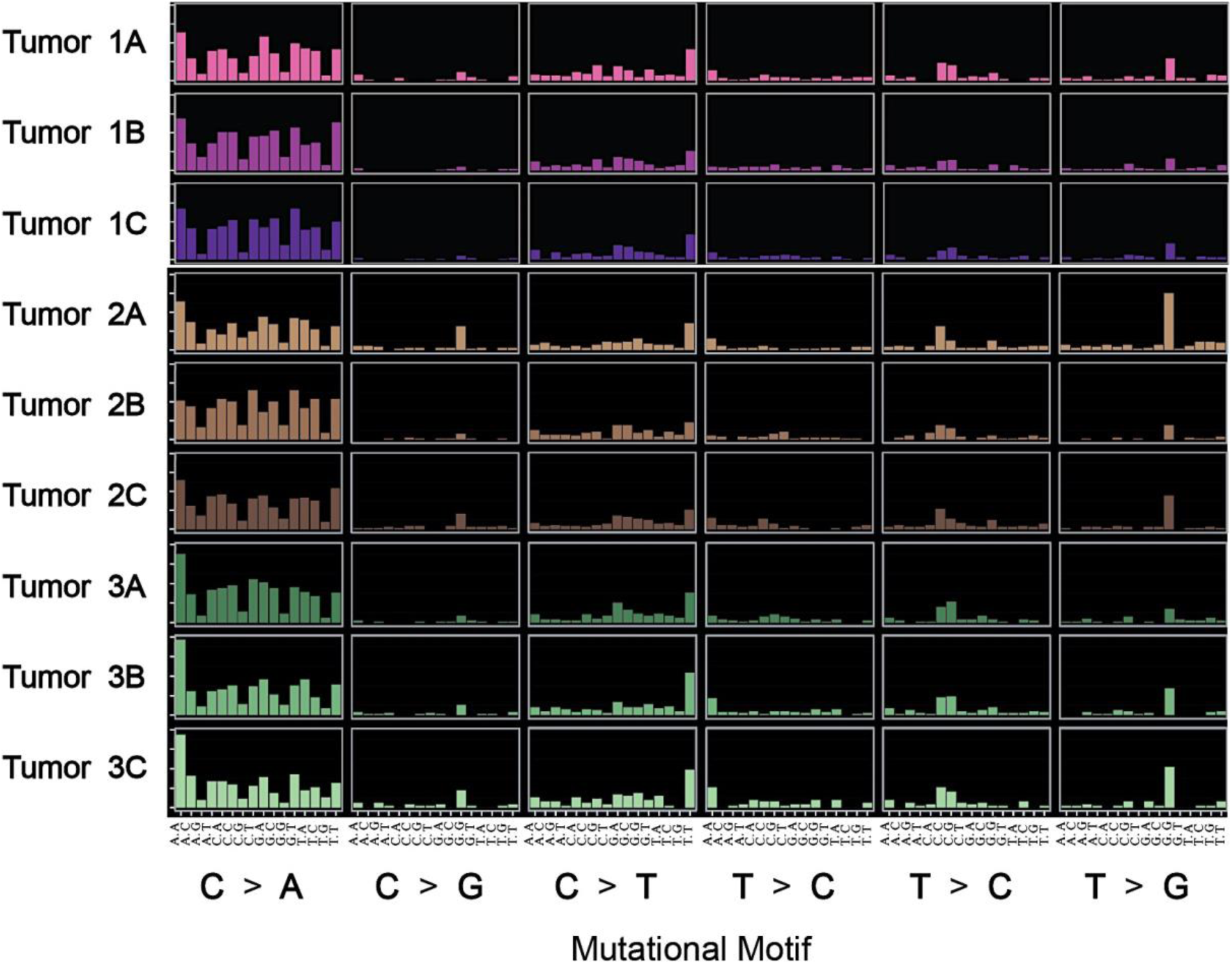
*Lrig1-Cre/+; Apcflox/+* colonic adenomas have a distinct pattern of somatic mutations dominated by C:G>A:T transversion point mutations. The somatic signature of mutations of the adenomas are plotted as rows; there are 9 adenomas from 3 mice. Each adenoma contains a nearly identical pattern of somatic mutations based on motifs and background sequence context. The somatic signature is characterized by an abundance of C to A transversion point mutations, which are caused by defects in repairing DNA adducts of guanine and are frequently observed in cancers associated with tobacco tar and *H. pylori* infections.

The tumors are highly heterogeneous in terms of the specific genetic loci mutated, displaying high genetic variability across tumors. Each adenoma contained a unique profile of ∼100 high-frequency and high-quality mutations predicted to cause significant functional disruptions based on amino acid conservation. Two genes were mutated in 4/9 tumors: *Muc4* and *Dhx8* (**Table S1**). Eleven genes were independently mutated in >30% of the tumors (**Figure S1**). The 40 most commonly mutated genes (>20%) are predicted to impact cell morphology and migration based of pathway analysis, and are present in over 10% of human CRC cases in The Cancer Genome Atlas (TCGA) database.

All of the tumors show some defects in DNA mismatch repair in both the mRNA and exomic DNA datasets. The specific genes affected are different for each of the tumors despite the remarkably similar mutational patterns. Mutated tumor suppressor genes, such as *Pi3kca, Nf1, Lrig1*, and DNA repair genes like *Msh4, Msh6*, and *Pold1* were observed. Unexpectedly, most tumors contained several additional mutations in canonical and non-canonical *Wnt* pathway genes.

In contrast to previous reports, a genetic loss of heterozygosity (LOH) of *Apc* was not observed in the tumor exome DNA (**Figure 3**). Allelic loss is typically considered to occur when the amount of tumor DNA is less than or equal to 50% of the value obtained from heterozygous tissue, due to the high risk of contamination from wildtype cells. However, in this case the tumor cells contained ∼50% of the adjacent normal DNA which is assumed to be homozygous for *Apc* since tamoxifen-induced Cre-recombination in the colonic epithelium of these mice is a very rare event. No spontaneous somatic mutations were identified in the *Apc* gene that were of meaningful quality and/or significance. No genetic copy number variations or insertions or deletions of *Apc*, or any other gene, were readily apparent in the data.

**Figure 3.**
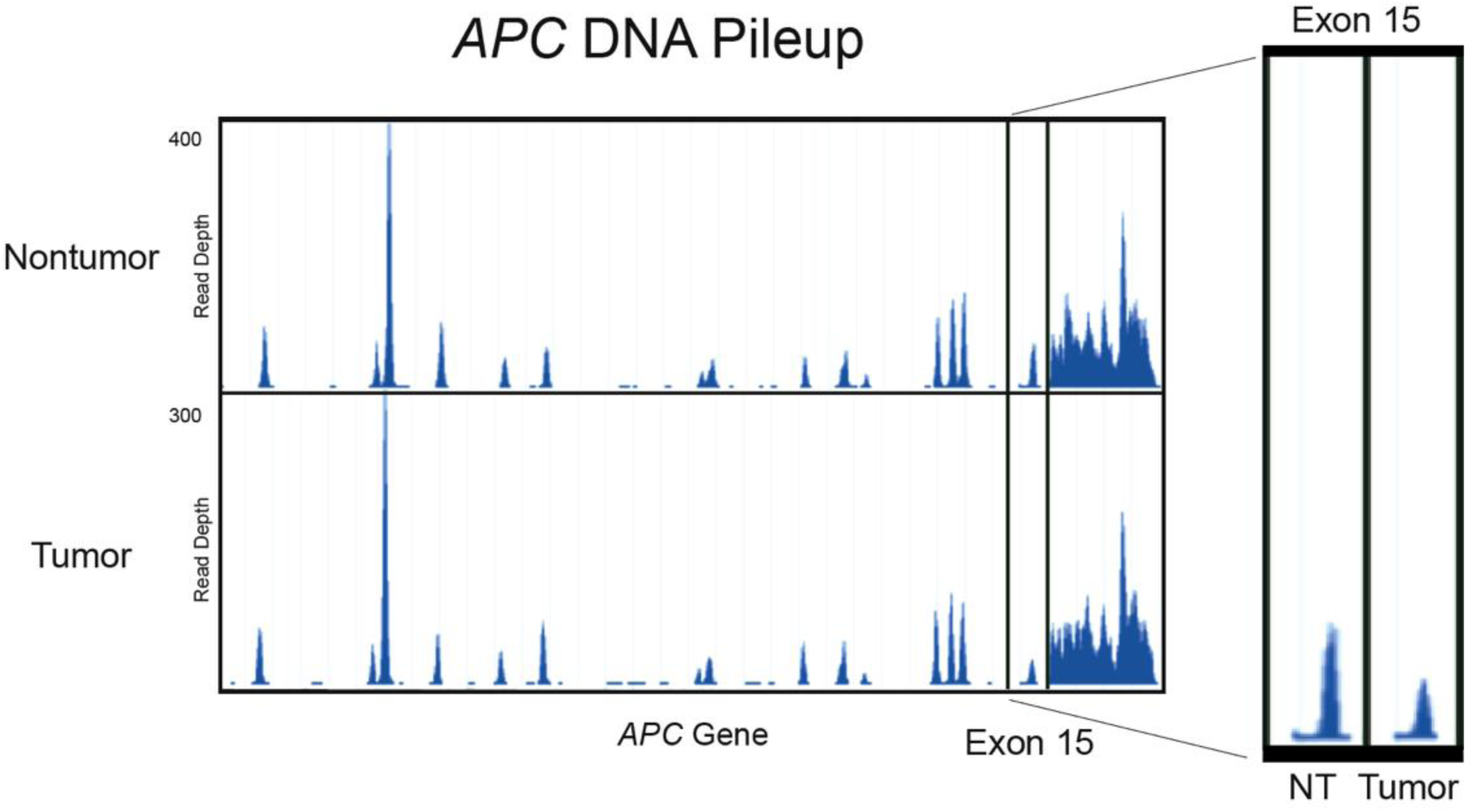
The loss of heterozygosity of the *Apc* gene was not observed through DNA sequencing. *Lrig1-Cre/+;Apcfl/+* adenomas are missing only one copy of exon 15 of the *Apc* gene. RNA sequencing reads from nontumor (top row) and adenoma (bottom row) are plotted as pooled pileups. Allelic loss is typically considered to occur when the amount of tumor DNA is less than or equal to 50% of the value obtained from heterozygous tissue, due to the high risk of contamination from wildtype cells. Here, the sequenced tumor cells contain ∼50% of the adjacent normal DNA which is assumed to be homozygous for wildtype *Apc.* No additional somatic mutations in *Apc* were observed across the nine tumors including genetic deletions, although many other *Wnt / B-catenin* genes were mutated.

Unlike the exomic DNA, the tumor mRNA displayed a clear lack of functional *Apc* across the entire dataset (n= 5/5 tumors). The adenomas express exclusively *Apc* transcripts lacking codon 580 in exon 15 (**Figure 4**), which is the same sequence as the original *Apc* transgene after tamoxifen-induced recombination has occurred. These results imply that the tumors have acquired the ability to selectively express the nonfunctional *Apc* transcripts. This epigenetic phenomenon appears to have occurred across all tumors, implying that the loss of heterozygosity of *Apc* was an early in the process of adenoma formation.

**Figure 4.**
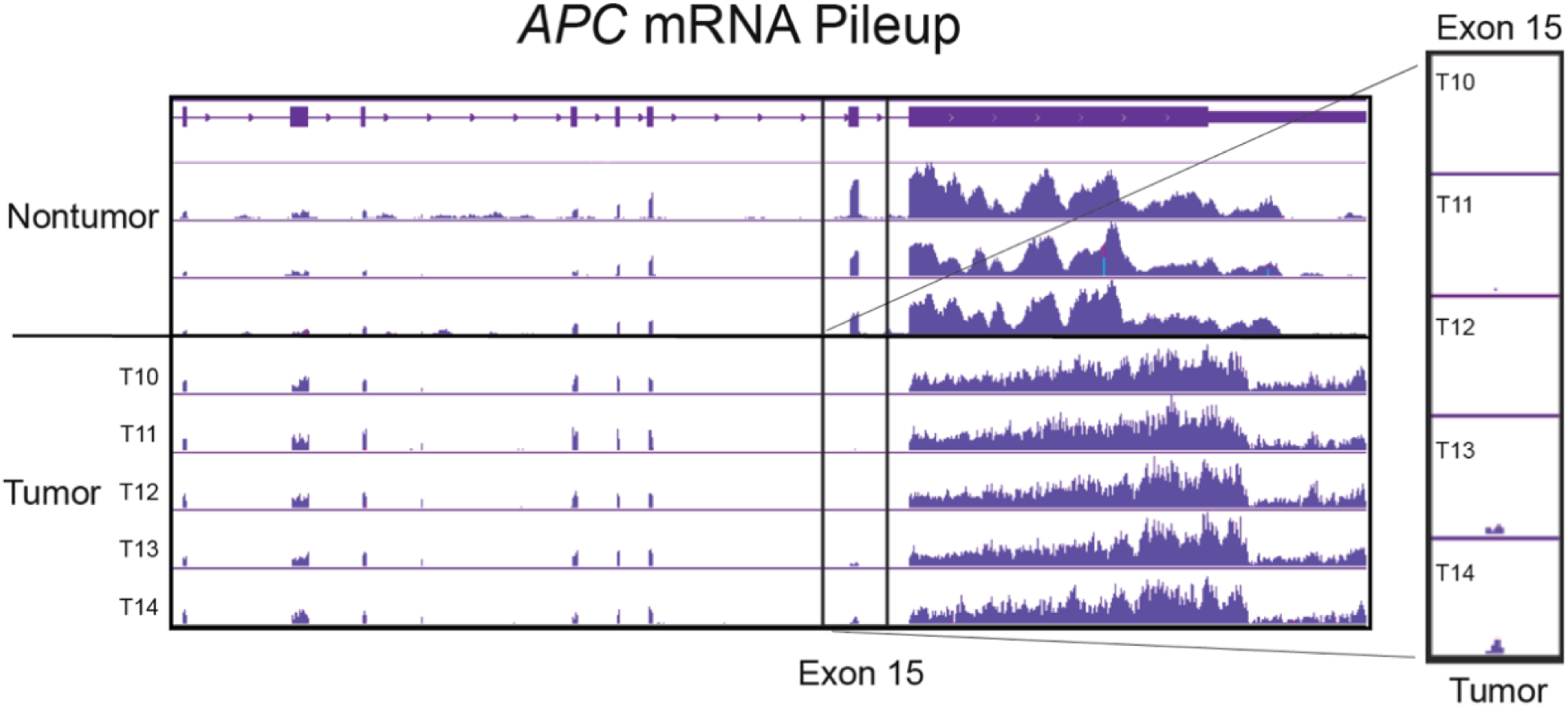
Lrig1-driven adenomas selectively express mutated *Apc* transcripts originating from a *Cre-*recombined transgene. The epigenetic loss of heterozygosity of *Apc* was apparent in the mRNA of *Lrig1-Cre/+;Apcfl/+* adenomas, which are missing exon 15 from *Apc* mRNA transcripts. RNA sequencing reads from nontumor (top three rows) and adenoma (bottom five rows) are plotted. The putative mechanism of *Apc* loss in these adenomas proceeds via epigenetic silencing of the wildtype allele and selective transcription of the mutant *Apc* gene, which lacks exon 15 after Cre-induced recombination occurs under the control of the *Lrig1* promoter. The purity of the tumor samples is demonstrated by the complete lack of *Apc* exon 15 from tumors T10, T11, and T12. Tumors T13 and T14, which have retained some expression of *Apc* exon 15, were found to have gene expression patterns most similar to wildtype cells based on RNA-Seq.

Despite their highly variable genomes, the tumors exhibited substantial transcriptome similarity to each other (**Figure S3**). Downregulation of the DNA repair genes *Msh3* and *Msh4* and several UDP glucuronosyltransferases was detected (**Table S2**). Upregulation of *Dkk* was also observed, highlighting the fact that tumor biomarkers can vary according to the stage, location, and genetic context of the tumor. Tumors consistently displayed abnormal patterns of intron retention in several RNA-binding genes (**Figure S4**). Compared to wildtype, the tumor transcripts display a loss of intron retention, which is a phenomenon sometimes observed in breast cancer ^30^. This observation suggests that modifications in post-transcriptional gene regulation are exploited by the tumor cells in order to gain additional selective advantages against the host.

The proposed mechanism of colonic adenoma formation in the *Lrig1-(Apc-flox)* model is the following: Epigenetic loss of *Apc* leads to a decrease in goblet cells and a corresponding increase in inflammatory reactive oxygen species (ROS) ^31^. Somatic mutations occur subsequently as a result of increased inflammation and provide the raw genetic material required for adenoma formation and progression. Natural selection of growth-promoting mutations leads to convergent evolution towards an invasive phenotype marked by invasion, increased survival, and alternate modes of *Wnt* pathway activation. The high incidence of adenomas in the distal colon can be explained by the higher prevalence of microbial-produced ROS localized to the distal colon and rectum.

## Discussion

In this study, the exomes and transcriptomes of conditionally induced murine colonic adenomas from *Lrig1-(Apc-flox)* mice were examined to assess the incidence and of genomic alterations during the early stage of tumorigenesis. Evidence of early genomic changes were apparent throughout the exomes and transcriptomes of the tumors sequenced in this study, including alterations in gene expression, splicing, and DNA repair. The tumors displayed a hypermutated phenotype with a high incidence of point mutation, which was quite unexpected due to the relatively low sequencing depth employed in this study and the early stage of the tumors. A high prevalence of genetic heterogeneity was observed in tumors from within the same mouse (intratumoral heterogeneity), as well as from sibling mice (intratumoral heterogeneity). Each tumor exome (n=9) containing a unique profile of ∼100 mutated alleles predicted as “high impact” on amino acid sequences based on the consensus of the two variant predictors: SNPeff and Ensembl’s VEP program.

Identifying somatic mutations from tumor genomic data is an inherently difficult and expensive technique due the high rate of false negative mutations and the inability of standard sequencing methods to detect low frequency alleles. The detection rate of false positive mutations in this study was significantly reduced with the use of high-stringency filtering steps. Matched nontumor samples are required to filter background germline variants from tumor genomic data, but even deep sequencing of a matched nontumor sample will not remove all false positive somatic mutations from a dataset. All animal and tumor genomic datasets have unique germline SNPs, repetitive, and noncoding regions that appear as somatic when using a standard reference genome for data alignment. The vast majority of these germline variants can be filtered out using the public dbSNP database, however a low level of germline SNPs will remain in due to dbSNP errors or insufficient coverage of that locus in the nontumor sample. Additional steps would be required to verify the somatic nature of mutations of interest, such as ultra-deep sequencing of the locus of original wild-type strain. The somatic mutations detected in this study are assumed real based on the clear signature of a pattern of C>A mutations, which is very unlikely to occur due to chance. To further support of the validity of the detected mutations, this dataset was compared to control nontumor samples and mouse glioblastoma whole genome DNA sequencing datasets (n=3) and no clear mutational patterns were observed in the controls. Anecdotally, the detected mutations were clearly malignant in nature but not so completely destructive as to kill the mice. Due to the low sequencing depth employed in this study, the somatic mutations detected likely represent only a fraction of the actual number present in the tumors.

The hypermutated phenotype of *Lrig1-(Apc-flox)* tumors could result from an increased rate of somatic mutations or a decreased rate of DNA mismatch repair. To identify potential driver mutational processes involved in *Lrig1-(Apc-flox)* tumorigenesis, the somatic signatures of mutation were determined based on mutation type, abundance, and background genomic sequence context. The adenomas have a distinct pattern of somatic mutations dominated by C:G>A:T transversion point mutations, which are often seen in smoking-induced lung cancers and stomach cancers caused by *H. pylori* infections ^11^. Glutathione synthases are typically responsible for removing oxidized carcinogenic adducts from guanine residues to prevent DNA mismatches; microbial-produced reactive oxygen species (ROS) are a likely source of guanine adducts. The high incidence of *Lrig1-(Apc-flox)* adenomas localized to the distal colon could be explained by the high levels of microbial-produced ROS inherent to the duodenum and rectum. In both *Gst-null, Apc*^*Min*^ and *Lrig1-(Apc-flox)* adenomas, the glucuronidation pathway is strongly downregulated and many inflammation markers were upregulated. It is unclear if the inflammation-associated gene expression changes are the cause or result of the increased mutation rate.

The *Lrig1-(Apc-flox*) model of CRC is notable for its high penetrance and rapid onset after the loss of just a single copy of *Apc*. Empirically observed rates of spontaneous loss of heterozygosity (LOH) in humans and mice tend to occur much more slowly, generally in the order of several months to years. For example, in human retinoblastoma tumors generally take years to develop in the retinas of children missing one copy of the tumor suppressor *Rb*, and human Familial adenomatous polyposis (FAP) intestinal tumors typically become malignant at around age 40. Murine cancer models of spontaneous tumor suppressor LOH (e.g. *p53, Brca1*) consistently produce tumors with much higher penetrance and growth rates than human LOH tumors. The rapid onset and extremely high tumor burden observed in *Lrig1-(Apc-flox*) mice are especially striking considering that prior to the spontaneous loss of an *Apc* allele, tamoxifen-induced Cre-recombinase expression under the control of the *Lrig1* promoter must first occur in the same cell as the random *Apc* mutation. In this study a genetic LOH of *Apc* was undetectable in the genomes of these tumors. This fact is unsurprising when one considers the extremely low theoretical likelihood of multiple spontaneous somatic LOH events at a single locus co-occurring with Cre-recombinase activation several times in one colon over the span of a few months. One simple explanation for the inability to detect a genetic LOH of *Apc* in these tumors is the high possibility of achieving false negative results due to contamination from stromal cells. It is worth noting that in the original report on these tumors a standard PCR genotyping assay was the method used to detect the loss of heterozygosity of *Apc*, which is an archaic and much less sensitive technique for genotyping compared to DNA sequencing. It is possible that preferential loss of the remaining wild-type *Apc* allele in the *Lrig1-(Apc-flox)* tumors is not a random event, but rather is directly engineered by the tumor via an unknown mechanism. Perhaps the double heterozygosity of *Lrig1* and *Apc* act synergistically with *Lrig1* haploinsufficiency to increase the possibility of an epigenetic LOH of *Apc*.

It is puzzling that most of the discovered somatic mutations in tumor suppressor genes are typically found in only a single gene copy. There are several possible explanations for this phenomenon, which are not mutually exclusive. One classical explanation is that mutations in genes from the same biochemical pathway are able to amplify the loss of one copy of a gene. Mutated regulatory sequences or epigenetic losses are another likely possibility. Haploinsufficiency of tumor suppressor genes may be a much more common mechanism of tumor growth than expected. Haploinsufficiency is currently an under-investigated topic due to the classical understanding of inherited tumor suppressors and the difficulty in achieving accurate SNP calls from heterogeneous tumors using next-generation sequencing. Perhaps the low sequencing depth used in this study is responsible for the assumed absence of the complementary tumor suppressor mutations, and this dataset may comprise a small fraction of the actual tumor mutation burden. Ultra-deep sequencing of the tumor and matched nontumor sample would provide additional needed insight into this issue.

Despite the normal appearance of the remaining *Apc* alleles in the tumor exome DNA, there was clear evidence of an epigenetic (‘*above the genes’*) LOH of exon 15 of *Apc* observed in the tumor mRNA. The assumed mechanism responsible for the lack of wildtype *Apc* expression is pre-transcriptional epigenetic silencing of the single remaining wildtype copy of *Apc* remaining after engineered transgene recombination. Unlike the tumor DNA, the *Apc* mRNA transcripts expressed by the tumor cells clearly display a copy number loss of exon 15. The expressed *Apc* transcripts contain an identical genetic sequence to that of the original *Apc* transgene following Cre-induced recombination, implying that the mutant transgene is being selectively expressed by the tumor cells through an epigenetic mechanism.

In order to develop therapeutic treatments that are less susceptible to drug resistance, it is important to understand the mechanisms underlying tumor formation and evolution. Countless genetic mechanisms regulating tumor growth have been elucidated and elegantly described in recent decades, but the cancer drugs designed based on this knowledge typically have had minimal effectiveness. The necessary steps required to consistently and reliably prevent CRC metastasis and drug resistance remain elusive and innovative approaches and avenues of research are critically required in order to cure CRC. Going forward, mutational processing mechanisms altered during early tumorigeneses make attractive therapeutic targets since they should theoretically be present in a large proportion of tumor cells and they are the drivers of tumor growth. Future studies are needed in order to understand the mechanisms underlying the processes of mutation generation used by tumors. One obvious follow-up experiment to this work would be to sequence *Gst-null; Apc*^*Min*^ tumors to assess the incidence of C > A point mutations and epigenetic losses of tumor suppressor genes including *Apc* and *Lrig1*.

The selection of advantageous mutations is a dynamic process, changing throughout the course of tumorigenesis. The results of this study demonstrate that the genomic events leading to tumor growth and invasion can happen early in tumorigenesis and that tumors can generate large pools of mutations early in the course of disease progression. The ability of a tumor to create and store mutations in a cache or reservoir until a later time provides an advantage to the tumor by optimizing its ability to dynamically modulate the selection of oncogenic mutations in a context-dependent manner throughout disease progression. Mutational cache creation provides tumors with the ability to conditionally express somatic mutations, ultimately leading to tumor metastasis and drug resistance. Understanding the role of mutation generation and storage during tumorigenesis may provide deep insights into the mechanism underlying the ability of cancer cells to modulate the gene expression of critical genes throughout tumorigenesis and metastasis, and could provide new avenues for more refined cancer treatment options.

## Materials & Methods

### Animals

Generation of *Lrigl<tml.l(cre/ERT)Rjc>* (*Lrig1-CreERT2, Apcflox*/+) mice was performed by the Coffey lab as previously described ^27^. The wildtype strain used for RNA-Seq was C57BL/6J. Mice were housed in a pathogen-free and light-controlled environment, fed standard chow, and allowed water ad libitum. Mice were given 2mg tamoxifen via i.p. injection for 3 consecutive days and were euthanized 80-120 days later. Procedures were approved by the Vanderbilt University Medical Center Animal Care and Use Program.

### Generation of Lrig1-driven colonic adenomas

Generation of *Lrig1-CreERT2, Apcflox*/+ tumors was performed by the Coffey lab as previously described ^32^. Briefly, adult (6-8-wk-old) *Lrig1-CreERT2/+;Apc-flox/+* mice were intraperitoneal-injected with 2mg tamoxifen (Sigma) in corn oil for 3 consecutive days. Multiple dysplastic duodenal colonic adenomas were observed 100 days later. Whole tumors (n=14) from 8 mice were excised and RNA (n=5 tumors) and DNA (n=9 tumors) were extracted ∼80-120 days following tamoxifen injection, depending on tumor burden. Tumor DNA and mRNA samples were shipped to HudsonAlpha Labs for genomic analysis during 2012-2015 (**Supplementary Table 3**).

### Messenger RNA and exomic DNA sequencing of Lrig1-driven colonic adenomas

Total RNA was extracted from tumors (n=5) and wildtype colon (n=1 pooled sample), as well as from wildtype *Lrig1*-expressing and *Lgr5-*expressing cells that were sorted and collected using flow cytometry and shipped frozen to HudsonAlpha Labs (Annie Powell, Coffey lab). Messenger RNA isolation, RNA-Seq library preparation, and cDNA sequencing were performed by HudsonAlpha Labs in 2012. The pooled wildtype colon sample was sequenced to a 5x greater depth than the individual tumor samples. Tumor DNA (n=9 tumors from n=3 mice) was extracted from isolated tumors consecutively with adjacent normal colon (n=3) and shipped frozen to HudsonAlpha Labs (Coffey lab). Tumor exome capturing, DNA-Seq library preparation, and sequencing were performed by HudsonAlpha Labs in 2015. Exomic DNA was sequenced to 12x average read depth.

### DNA read alignment and processing

Raw fastq reads were cleaned to remove low quality bases at the ends of the reads using Stacks (v 1.35) process_shortreads ^33^. Cleaned reads were aligned to the *mus musculus* (house mouse) genome (2011 assembly, UCSC Genome Browser assembly ID mm10, Genome Reference Consortium Mouse Build 38, Accession GCA_000001635.2) using Bowtie2 (v2.2.1), in default-sensitive mode ^34^. Sam/bam files were sorted and indexed using SAMtools (v0.1.18) ^35^ and Picard Tools (v1.92) (Broad Institute, MIT License). Base quality score recalibration and indel realignment were performed according to GATK best practices ^36^.

### Mutation calling, filtering, and visualization

Somatic mutations were called using SeuratSomatic (v2.5), a GATK module ^37-39^. Somatic variants were called using a minimum variant coverage of 4 reads and minimum quality score of Q10. To ensure that the identified mutations were somatic in nature, matched normal tissue samples were used to filter germline variants. Residual germline polymorphisms were removed by filtering all germline variants reported on NCBI’s dbSNP database ^40^. To reduce the occurrence of sequencing errors in the data, variants were called only if they were present in both the forward and reverse DNA strand ^41^. To confirm that the discovered mutations were not due to alignment artifacts, the presence of representative variants was verified in the raw sequencing reads using grep and BLAST. The predicted effect of the mutations on protein function was determined with Ensembl’s variant effect predictor (VEP) program ^42^ as well as the SnpEff variant annotation and effect prediction tool ^43^. The somatic signature of the mutations was identified using the R package SomaticSignatures ^39^. Genomic data was visualized using the R packages GenVisR and ggplot2 ^44,45^ Negative control nontumor samples and mouse glioblastoma whole genome DNA sequencing datasets (n=3, provided by Hui Zong Lab) were compared to this dataset.

### RNA sequencing and analysis

Whole tumors (n=6) as well as normal wildtype colon (n=3), were excised and mRNA was sequenced to 25x average read depth. Cleaned cDNA reads were aligned to the mouse mm10 genome (Accession GCA_000001635.2) with STAR(v1.0). Changes in gene expression were identified using the R package DESeq2 and visualized with the R packages DESeq2, ggplot2, and Pathview. Custom bigwig tracks were created on the UCSC Genome browser in order to assess copy number variations ^46^.

## Supplementary Figures

**Figure S1.** Putative Driver Mutations in *Lrig1-(Apc-flox)* Colonic Adenomas.

**Figure S2A.**MAplot differentially-expressed genes in *Lrig1-(Apc-flox)* colonic adenomas.

**Figure S2B.** Heatmap visualization of highly significant differentially-expressed genes in *Lrig1-(Apc-flox)* colonic adenomas.

**Figure S3.** Principal components analysis of tumor transcriptome data compared to wildtype colon.

**Figure S4**. Lrig1-Cre/+;Apcfl/+ colonic adenomas have abnormal splicing patterns.

## Supplementary Tables

**Table S1**. Somatic mutations identified in Lrig1-Cre/+;Apcfl/+ colonic adenomas

**Table S2**. RNA-Seq results for in Lrig1-Cre/+;Apcfl/+ colonic adenomas vs. wildtype colon

**Table S3**. Mouse Sample IDs

## Acknowledgements

Bob Coffey conceived of the original idea for creating and sequencing the *Lrig1-(Apc-flox)* tumors and provided the sequencing data used for this project. Annie Zemper acquired funding and provided general laboratory supervision, administrative and technical support, writing assistance, and helpful discussions. HudsonAlpha Genomic Services Lab in Huntsville, AL performed all genomic library preparations and sequencing. Nicholas Stiffler and Doug Turnbull provided bioinformatics support. Funding for this research was provided by the University of Oregon and the following sources: NIH T32 CA119925 (Vanderbilt University Medical Center, Integrated Biological Systems Training in Oncology), NIH R25 CA136440 (Vanderbilt University Medical Center, Training in Cellular and Molecular Imaging of Cancer), NCI R01 CA151566 (Vanderbilt University Medical Center, Cell-of-origin effects on development of colon cancer). The results published here are in part based upon data generated by the TCGA Research Network: https://www.cancer.gov/tcga. This work benefited from access to the University of Oregon’s high performance computer, Talapas.

## References

1. Nitsche, U. et al. Meta-analysis of outcomes following resection of the primary tumour in patients presenting with metastatic colorectal cancer. Br J Surg 105, 784–796, doi:10.1002/bjs.10682 (2018).

2. Advani, S. & Kopetz, S. Ongoing and future directions in the management of metastatic colorectal cancer: Update on clinical trials. J Surg Oncol 119, 642–652, doi:10.1002/jso.25441 (2019).

3. Sasaki, N. & Clevers, H. Studying cellular heterogeneity and drug sensitivity in colorectal cancer using organoid technology. Curr Opin Genet Dev 52, 117–122, doi:10.1016/j.gde.2018.09.001 (2018).

4. Roerink, S. F. et al. Intra-tumour diversification in colorectal cancer at the single-cell level. Nature 556, 457–462, doi:10.1038/s41586-018-0024-3 (2018).

5. Linnekamp, J. F. et al. Consensus molecular subtypes of colorectal cancer are recapitulated in in vitro and in vivo models. Cell Death Differ 25, 616–633, doi:10.1038/s41418-017-0011-5 (2018).

6. Albuquerque, C. et al. The ‘just-right’ signaling model: APC somatic mutations are selected based on a specific level of activation of the beta-catenin signaling cascade. Hum Mol Genet 11, 1549–1560, doi:10.1093/hmg/11.13.1549 (2002).

7. Smits, R. et al. Somatic Apc mutations are selected upon their capacity to inactivate the beta-catenin downregulating activity. Genes Chromosomes Cancer 29, 229–239 (2000).

8. Christie, M. et al. Different APC genotypes in proximal and distal sporadic colorectal cancers suggest distinct WNT/beta-catenin signalling thresholds for tumourigenesis. Oncogene 32, 4675–4682, doi:10.1038/onc.2012.486 (2013).

9. Fearon, E. R. & Vogelstein, B. A genetic model for colorectal tumorigenesis. Cell 61, 759–767 (1990).

10. Alexandrov, L. B., Nik-Zainal, S., Wedge, D. C., Campbell, P. J. & Stratton, M. R. Deciphering signatures of mutational processes operative in human cancer. Cell Rep 3, 246–259, doi:10.1016/j.celrep.2012.12.008 (2013).

11. Alexandrov, L. B. et al. Signatures of mutational processes in human cancer. Nature 500, 415–421, doi:10.1038/nature12477 (2013).

12. Alexandrov, L. B. & Stratton, M. R. Mutational signatures: the patterns of somatic mutations hidden in cancer genomes. Curr Opin Genet Dev 24, 52–60, doi:10.1016/j.gde.2013.11.014 (2014).

13. Nadkarni, A. et al. Nucleotide Excision Repair and Transcription-coupled DNA Repair Abrogate the Impact of DNA Damage on Transcription. J Biol Chem 291, 848–861, doi:10.1074/jbc.M115.685271 (2016).

14. Diouf, B. et al. Somatic deletions of genes regulating MSH2 protein stability cause DNA mismatch repair deficiency and drug resistance in human leukemia cells. Nat Med 17, 1298–1303, doi:10.1038/nm.2430 (2011).

15. Woerner, S. M. et al. Detection of coding microsatellite frameshift mutations in DNA mismatch repair-deficient mouse intestinal tumors. Mol Carcinog 54, 1376–1386, doi:10.1002/mc.22213 (2015).

16. Cabelof, D. C. Haploinsufficiency in mouse models of DNA repair deficiency: modifiers of penetrance. Cell Mol Life Sci 69, 727–740, doi:10.1007/s00018-011-0839-7 (2012).

17. Wong, V. W. et al. Lrig1 controls intestinal stem-cell homeostasis by negative regulation of ErbB signalling. Nat Cell Biol 14, 401–408, doi:10.1038/ncb2464 (2012).

18. Faraz, M., Herdenberg, C., Holmlund, C., Henriksson, R. & Hedman, H. A protein interaction network centered on leucine-rich repeats and immunoglobulin-like domains 1 (LRIG1) regulates growth factor receptors. J Biol Chem 293, 3421–3435, doi:10.1074/jbc.M117.807487 (2018).

19. Neirinckx, V., Hedman, H. & Niclou, S. P. Harnessing LRIG1-mediated inhibition of receptor tyrosine kinases for cancer therapy. Biochim Biophys Acta Rev Cancer 1868, 109–116, doi:10.1016/j.bbcan.2017.02.007 (2017).

20. Munoz, J. et al. The Lgr5 intestinal stem cell signature: robust expression of proposed quiescent ‘+4’ cell markers. EMBO J 31, 3079–3091, doi:10.1038/emboj.2012.166 (2012).

21. Yu, S. et al. Expression of LRIG1, a Negative Regulator of EGFR, Is Dynamically Altered during Different Stages of Gastric Carcinogenesis. Am J Pathol 188, 2912–2923, doi:10.1016/j.ajpath.2018.08.006 (2018).

22. Torigoe, H. et al. Tumor-suppressive effect of LRIG1, a negative regulator of ErbB, in non-small cell lung cancer harboring mutant EGFR. Carcinogenesis 39, 719–727, doi:10.1093/carcin/bgy044 (2018).

23. Li, W. & Zhou, Y. LRIG1 acts as a critical regulator of melanoma cell invasion, migration, and vasculogenic mimicry upon hypoxia by regulating EGFR/ERK-triggered epithelial-mesenchymal transition. Biosci Rep 39, doi:10.1042/BSR20181165 (2019).

24. Fontao, F., Barnes, L., Kaya, G., Saurat, J. H. & Sorg, O. From the Cover: High Susceptibility of Lrig1 Sebaceous Stem Cells to TCDD in Mice. Toxicol Sci 160, 230–243, doi:10.1093/toxsci/kfx179 (2017).

25. Lindquist, D. et al. LRIG1 negatively regulates RET mutants and is downregulated in thyroid cancer. Int J Oncol 52, 1189–1197, doi:10.3892/ijo.2018.4273 (2018).

26. Mao, F. et al. Lrig1 is a haploinsufficient tumor suppressor gene in malignant glioma. Oncogenesis 7, 13, doi:10.1038/s41389-017-0012-8 (2018).

27. Powell, A. E. et al. The pan-ErbB negative regulator Lrig1 is an intestinal stem cell marker that functions as a tumor suppressor. Cell 149, 146–158, doi:10.1016/j.cell.2012.02.042 (2012).

28. Chatterjee, A. & Gupta, S. The multifaceted role of glutathione S-transferases in cancer. Cancer Lett 433, 33–42, doi:10.1016/j.canlet.2018.06.028 (2018).

29. Ritchie, K. J., Walsh, S., Sansom, O. J., Henderson, C. J. & Wolf, C. R. Markedly enhanced colon tumorigenesis in Apc(Min) mice lacking glutathione S-transferase Pi. Proc Natl Acad Sci U S A 106, 20859–20864, doi:10.1073/pnas.0911351106 (2009).

30. Dvinge, H. & Bradley, R. K. Widespread intron retention diversifies most cancer transcriptomes. Genome Med 7, 45, doi:10.1186/s13073-015-0168-9 (2015).

31. Yang, K. et al. Interaction of Muc2 and Apc on Wnt signaling and in intestinal tumorigenesis: potential role of chronic inflammation. Cancer Res 68, 7313–7322, doi:10.1158/0008-5472.CAN-08-0598 (2008).

32. Powell, A. E. et al. Inducible loss of one Apc allele in Lrig1-expressing progenitor cells results in multiple distal colonic tumors with features of familial adenomatous polyposis. Am J Physiol Gastrointest Liver Physiol 307, G16–23, doi:10.1152/ajpgi.00358.2013 (2014).

33. Catchen, J., Hohenlohe, P. A., Bassham, S., Amores, A. & Cresko, W. A. Stacks: an analysis tool set for population genomics. Mol Ecol 22, 3124–3140, doi:10.1111/mec.12354 (2013).

34. Langmead, B., Trapnell, C., Pop, M. & Salzberg, S. L. Ultrafast and memory-efficient alignment of short DNA sequences to the human genome. Genome Biol 10, R25, doi:10.1186/gb-2009-10-3-r25 (2009).

35. Li, H. et al. The Sequence Alignment/Map format and SAMtools. Bioinformatics 25, 2078–2079, doi:10.1093/bioinformatics/btp352 (2009).

36. Van der Auwera, G. A. et al. From FastQ data to high confidence variant calls: the Genome Analysis Toolkit best practices pipeline. Curr Protoc Bioinformatics 43, 11 10 11–33, doi:10.1002/0471250953.bi1110s43 (2013).

37. Christoforides, A. et al. Identification of somatic mutations in cancer through Bayesian-based analysis of sequenced genome pairs. BMC Genomics 14, 302, doi:10.1186/1471-2164-14-302 (2013).

38. McKenna, A. et al. The Genome Analysis Toolkit: a MapReduce framework for analyzing next-generation DNA sequencing data. Genome Res 20, 1297–1303, doi:10.1101/gr.107524.110 (2010).

39. Gehring, J. S., Fischer, B., Lawrence, M. & Huber, W. SomaticSignatures: inferring mutational signatures from single-nucleotide variants. Bioinformatics 31, 3673–3675, doi:10.1093/bioinformatics/btv408 (2015).

40. Sherry, S. T. et al. dbSNP: the NCBI database of genetic variation. Nucleic Acids Res 29, 308–311, doi:10.1093/nar/29.1.308 (2001).

41. Preston, J. L. et al. High-specificity detection of rare alleles with Paired-End Low Error Sequencing (PELE-Seq). BMC Genomics 17, 464, doi:10.1186/s12864-016-2669-3 (2016).

42. McLaren, W. et al. The Ensembl Variant Effect Predictor. Genome Biol 17, 122, doi:10.1186/s13059-016-0974-4 (2016).

43. Cingolani, P. et al. A program for annotating and predicting the effects of single nucleotide polymorphisms, SnpEff: SNPs in the genome of Drosophila melanogaster strain w1118; iso-2; iso-3. Fly (Austin) 6, 80–92, doi:10.4161/fly.19695 (2012).

44. Skidmore, Z. L. et al. GenVisR: Genomic Visualizations in R. Bioinformatics 32, 3012–3014, doi:10.1093/bioinformatics/btw325 (2016).

45. Ginestet, C. ggplot2: Elegant Graphics for Data Analysis. J R Stat Soc a Stat 174, 245–245, doi:DOI 10.1111/j.1467-985X.2010.00676_9.x (2011).

46. Raney, B. J. et al. Track data hubs enable visualization of user-defined genome-wide annotations on the UCSC Genome Browser. Bioinformatics 30, 1003–1005, doi:10.1093/bioinformatics/btt637 (2014).

