## Supplementary Figures for "Epigenetic loss of heterozygosity of *Apc* and an inflammation-associated mutational signature detected in *Lrig1*^+/-^-driven murine colonic adenomas"

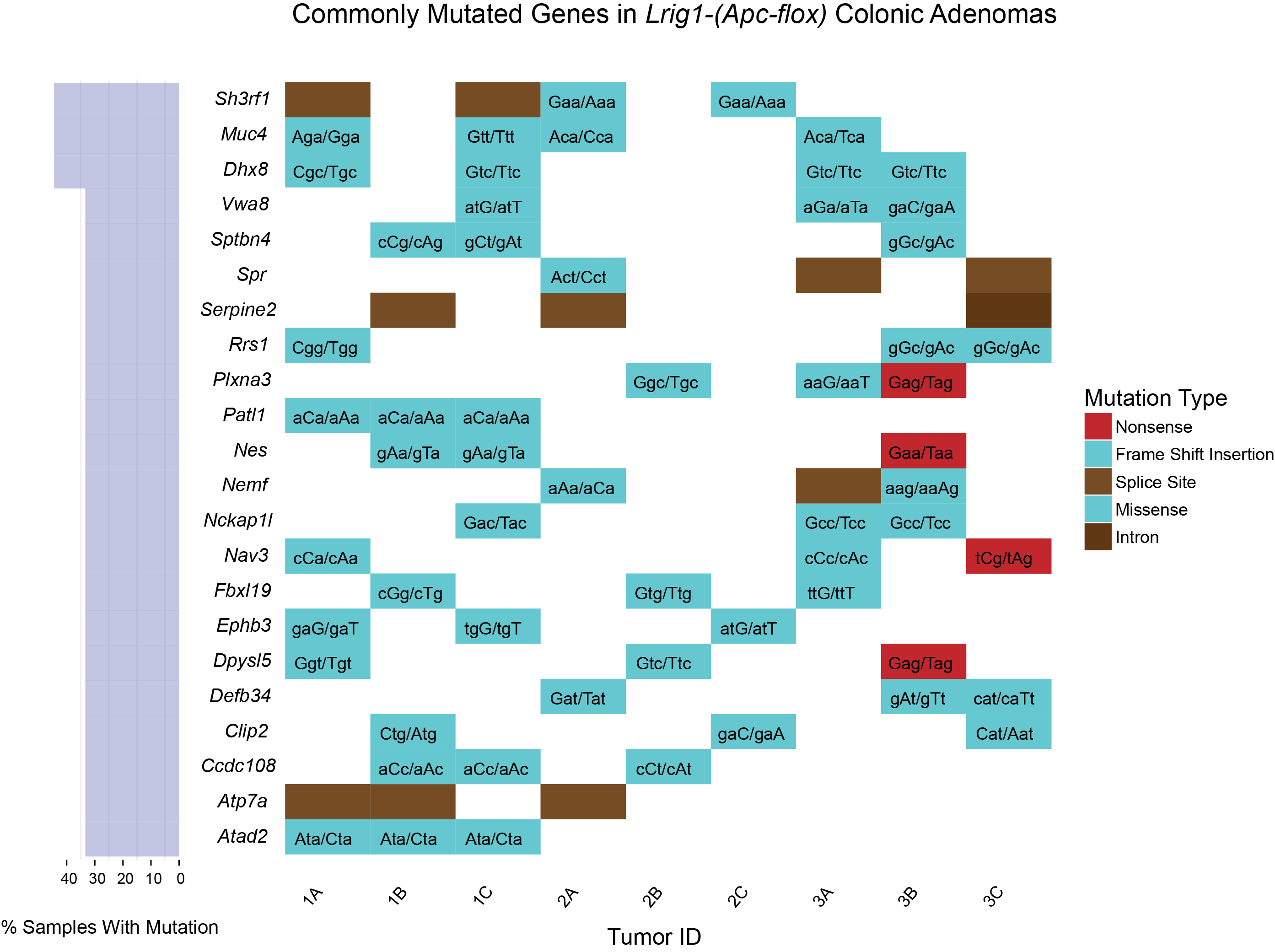

**Figure S1. Putative Driver Mutations in *Lrig1-(Apc-flox)* Colonic Adenomas**. Eleven genes were independently mutated in >30% of the tumors; these are shown as rows. Each column represents a single colonic tumor. The majority of the mutations are missense, C:G>A:T transversions. This plot was created using GenVisR.

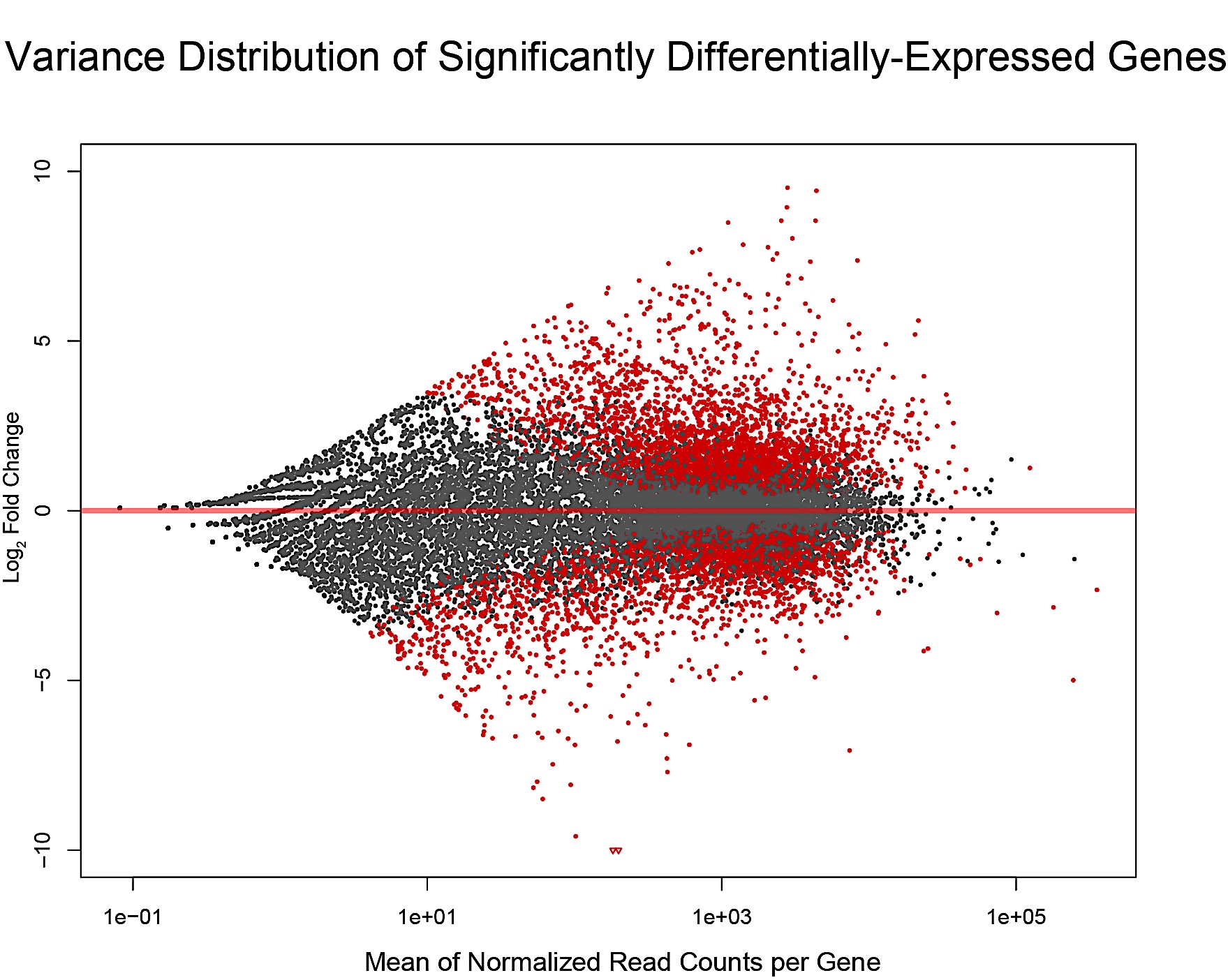

**Figure S2A.MAplot differentially-expressed genes in *Lrig1-(Apc-flox)* colonic adenomas.** Each dot represents a gene; genes in red are significantly different in tumor vs. wildtype mRNA. Genes above the x-axis are upregulated in tumors and genes below the x-axis are downregulated. Genes at the peaks of the pattern are highlighted in Figure S2B. This plot was created using DESeq2.

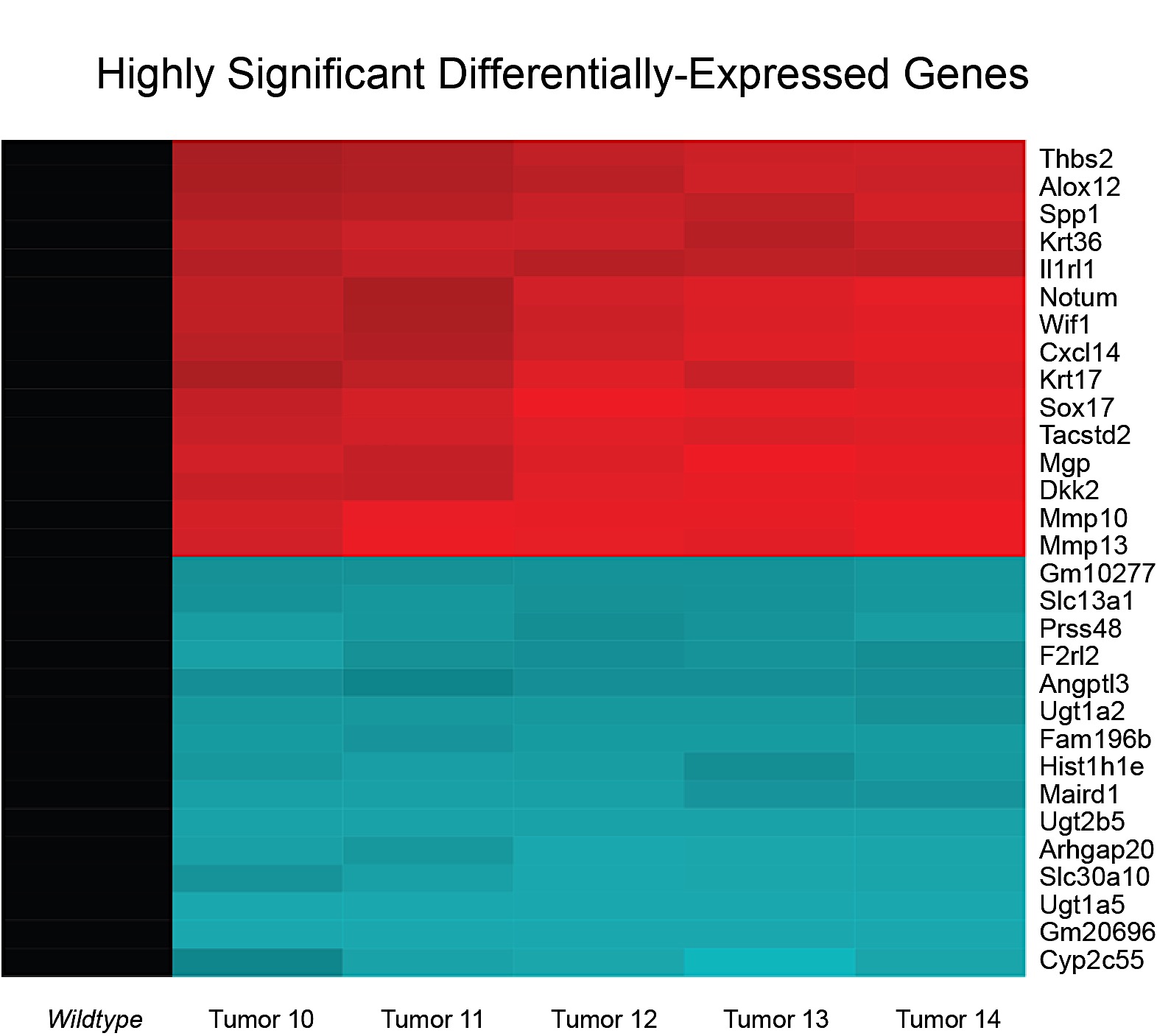

**Figure S2B. Heatmap visualization of highly significant differentially-expressed genes in *Lrig1-(Apc-flox)* colonic adenomas.** Rows are genes and colums are tumors. Genes colored red are upregulated (top 15 rows) and genes colored blue are downregulated (bottom 15 rows. Expression levels were normalized against the wildtype colon sample (black column on left). Many downregulated genes are involved in detoxification of exogenous carcinogens. This plot was created using ggplot2

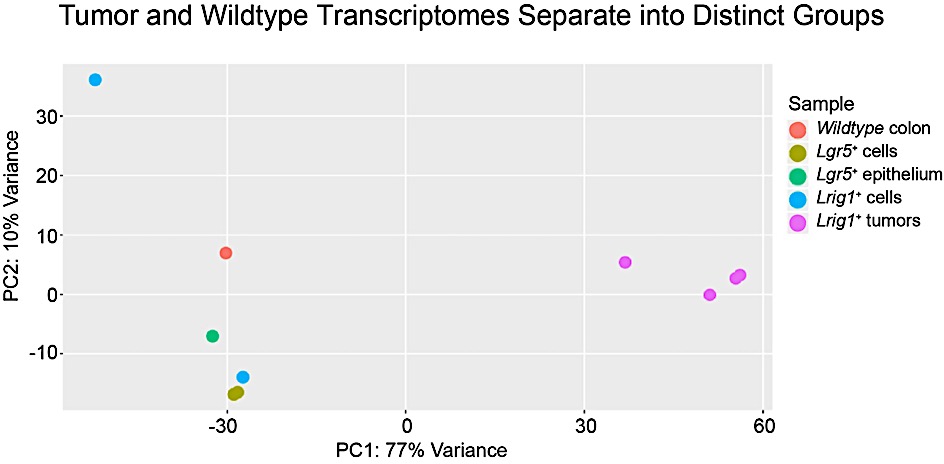

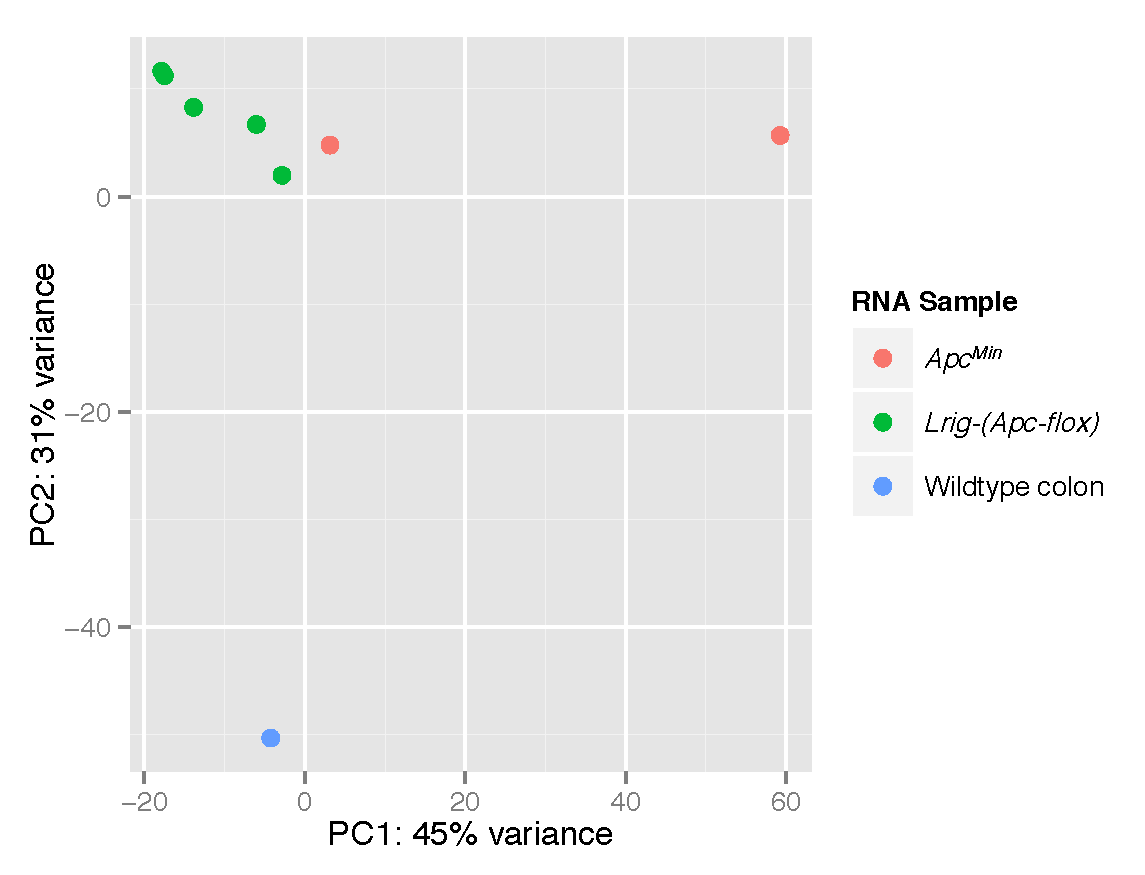

**Figure S3. Principal components analysis of tumor transcriptome data compared to wildtype colon.** Tumor and wildtype colon transcriptomes separate into distinct groups based on principal components analysis. This plot was created using DESeq2.

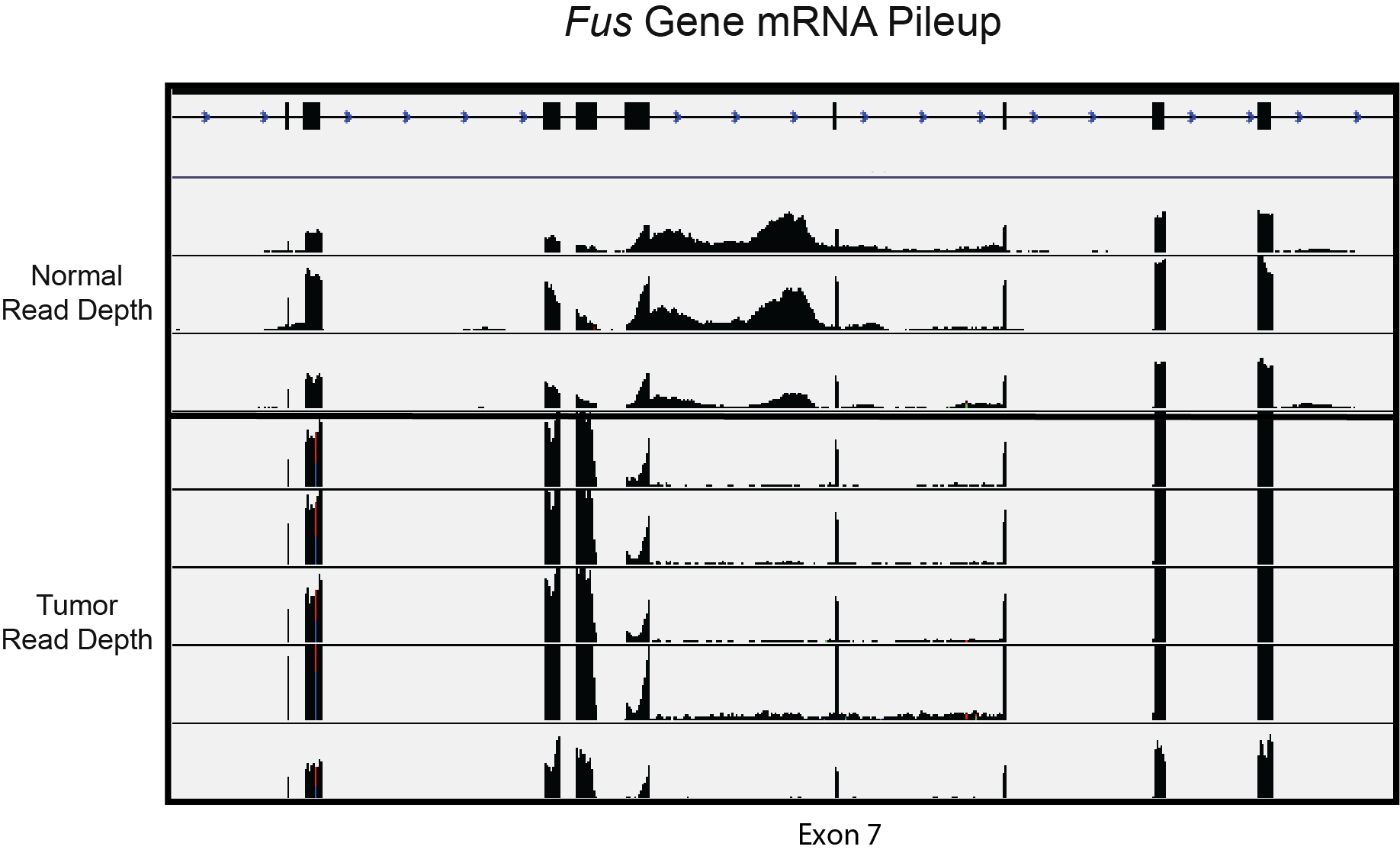

**Figure S4A. Lrig1-Cre/+;Apcfl/+ colonic adenomas have abnormal splicing patterns.** The Lrig1-Cre/+;Apcfl/+ tumor transcriptome is characterized by a loss of intron retention in RNA-binding genes. Abnormal patterns of gene splicing were observed throughout the mRNA pileups of genes involved in RNA-binding, including the gene Fus. RNA sequencing reads from nontumor (top) and adenoma (bottom) are plotted. Intron retention is a common feature of tumor genomes, but intron loss has been observed in breast cancer (Dvinge et al. Genome Med 2015). This plot was created using the UCSC genome browser.

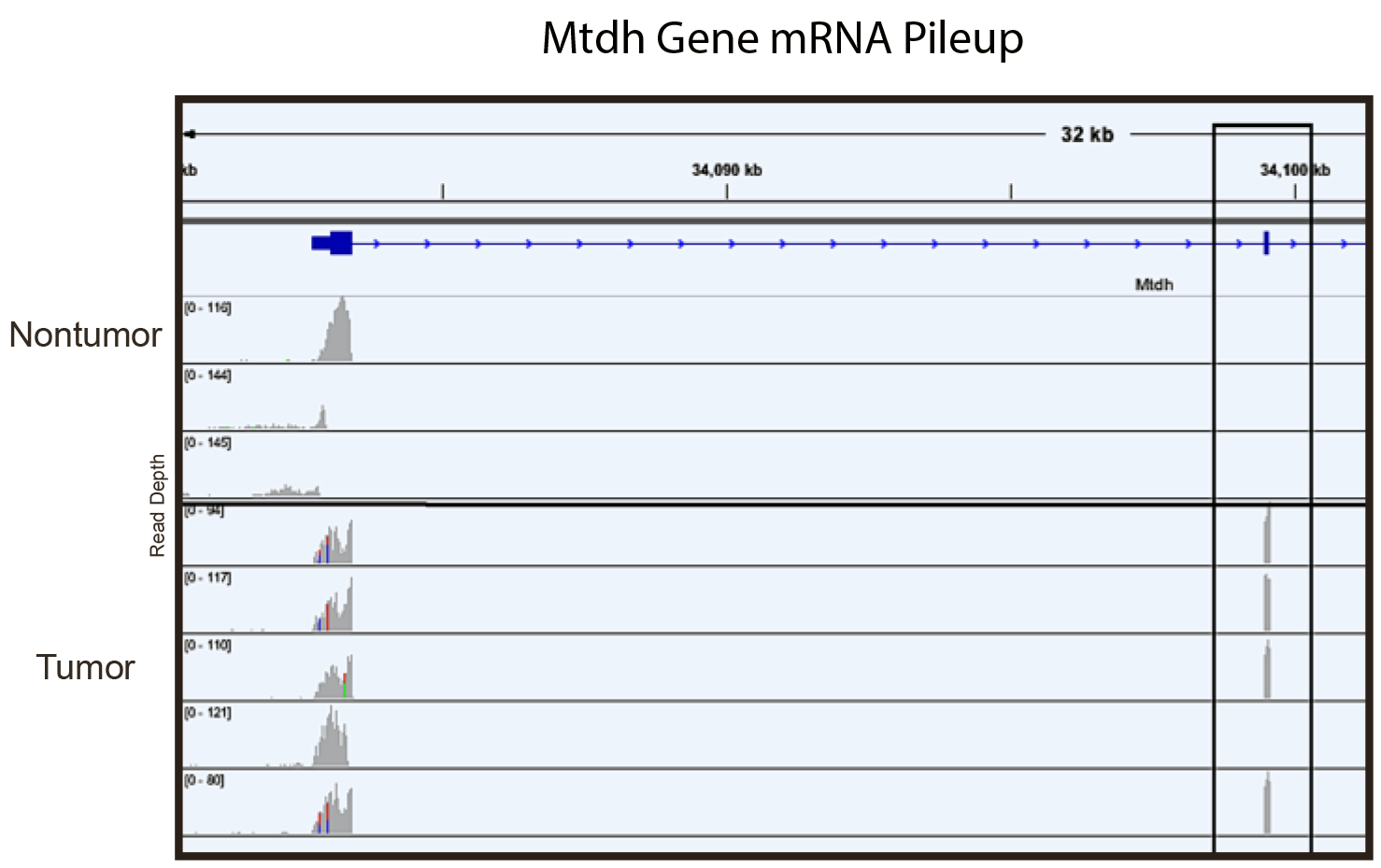

**Figure S4B. Lrig1-Cre/+;Apcfl/+ colonic adenomas have abnormal splicing patterns.** Abnormal patterns of gene splicing were observed in the CRC-associated gene *Mtdh* This plot was created using the UCSC genome browser.
